# Single-molecule imaging reveals activity-dependent regulation of *Camk2a* mRNAs at dendritic spines

**DOI:** 10.1101/2025.09.01.673561

**Authors:** Dong-Woo Hwang, Karthik Krishnamurthy, Arya Nagare, Robert H. Singer, Sulagna Das

## Abstract

Postsynaptic calcium/calmodulin-dependent protein kinase type II (CaMKII) integrates fleeting Ca^2+^ transients into long-term synaptic potentiation (LTP). A persistent presence of CaMKIIα at dendritic spines during the maintenance of LTP facilitates the prolongation of synaptic transmission. Yet, it remains unclear how the perpetuation of CaMKIIα, despite protein turnover, is achieved at dendritic spines. By visualizing endogenous *Camk2a* mRNAs at single molecule resolution using a newly developed mouse model, we identified a rapid activity-dependent localization of mRNAs to stimulated spines near the postsynaptic density (PSD) of hippocampal neurons. This spine localization was conferred by *cis*-acting regulatory elements termed cytoplasmic polyadenylation elements (CPEs) in *Camk2a* mRNA. Spine-localized *Camk2a* underwent on-site translation, which persisted for extended periods. These findings uncovered a novel local regulation of *Camk2a* mRNA, which serves to supply dendritic spines with a steady pool of highly concentrated CaMKIIα for maintaining long-lasting synaptic plasticity.

## Introduction

CaMKII is known to integrate a transient Ca^2+^ influx, established through the postsynaptic N-methyl-D-aspartate receptors (NMDARs), into downstream signaling cascades that underlie long-lasting plasticity at dendritic spines of excitatory neurons (*1–4*). The α subunit (i.e. CaMKIIα) predominantly constitutes dodecameric CaMKII holoenzymes that are associated with the PSD (*5–7*). The postsynaptic CaMKIIα is shown to mediate LTP, which is the cellular correlate for learning, after the initial Ca^2+^ signal rapidly wanes (*1, 2*). Mounting evidence has attributed this regulatory role of CaMKIIα in LTP to Ca^2+^-independent functions of the protein, such as autonomous phosphorylation at the critical regulatory residue Threonine 286 and its interaction with the NMDAR subunit GluN2B.

Studies of subcellular distribution of CaMKIIα in rodent hippocampal neurons have uncovered swift redistribution of the protein at dendritic spines in response to activation of the NMDARs, which persists over minutes and hours (*8–11*). The timing of this postsynaptic accumulation of the protein overlaps with the maintenance phases of various forms of long-lasting synaptic plasticity, including structural long-term synaptic plasticity (sLTP) (*12, 13*). As such, CaMKIIα has been postulated as a molecule that serves to perpetuate synaptic transmission at a particular synapse by exerting its Ca^2+^-independent functions (*9–11*). However, it has yet to be completely understood how CaMKIIα remains persistently concentrated in a restricted spine for extended periods against dynamic protein turnover, prompted by diffusion and degradation (*2, 14, 15*).

Biochemical studies have found *Camk2a* mRNAs in the synaptosomes, isolated from the resting rat visual cortex and mouse brain, raising the possibility of local regulation such as protein synthesis at synapses (*16–19*). Indeed, stimulation of the respective synaptosomes by a brief light exposure of dark-reared animals (i.e. visual experience) and activation of the NMDARs (i.e. Ca^2+^ influx) leads to increases in CaMKIIα protein levels, supporting activity-dependent protein synthesis at synapses. Importantly, disruption of dendritic localization of *Camk2a* by genetic deletion of the 3’untranslated regions (UTR) in a mouse model showed a failure to maintain the late phases of LTP despite the presence of residual CaMKIIα at the PSD, sourced from a somatic pool (*20*). These animals with impaired *Camk2a* mRNA localization are deficient in various forms of learning and memory, establishing the importance of RNA localization in driving behavioral outcomes. These findings provide evidence that activity-dependent synthesis of CaMKIIα at synapses is indispensable for the maintenance of long-term synaptic plasticity. Importantly, the synaptic protein synthesis is a strong suggestive of a self-perpetuating mechanism that is maintained by newly synthesized CaMKIIα.

Single-molecule imaging of the endogenous synaptic mRNAs, encoding protein components of synapses, has revealed that neurons harness various modes of local regulation to establish a local supply of synaptic proteins near a stimulated spine where the initial Ca^2+^ signal is elicited (*21–23*). Here, we report unique local regulation of *Camk2a* mRNA in mouse hippocampal neurons. This regulation is featured by unprecedented time and spatial scales, in which the mRNAs localized to stimulated dendritic spines near the PSD in a few minutes upon stimulation; CPEs in the 3’UTR of *Camk2a* were necessary and sufficient for this activity-dependent spine localization; localized-*Camk2a* remained *in situ* to undergo persistent on-site protein synthesis for up to 60 minutes. We propose this novel local regulation of *Camk2a* mRNA as a synaptic mechanism that serves to establish an enduring pool of highly concentrated CaMKIIα in a spatially restricted spine, wherein a sheer protein abundance would drive incorporation of the newly made α subunits that bear a temporal record of synaptic activation into the pre-existing dodecameric CaMKII holoenzymes. Therefore, this mechanism provides a long-term basis for perpetuating synaptic plasticity.

## Results

### Generation of *Camk2a* stem-loop knock-in mouse model and the detection of the *Camk2a* mRNA at activated postsynaptic dendritic spines in hippocampal neurons

*Camk2a* mRNAs is one of the most abundant mRNAs enriched in the neuropil of murine hippocampus, where it presumably undergoes dendritic and synaptic translation for long term synaptic plasticity (*17, 18, 24, 25*). The spatiotemporal distribution of the endogenous *Camk2a* mRNA with respect to the activated postsynaptic dendritic spines has yet to be thoroughly investigated. To directly probe activity-dependent postsynaptic localization of *Camk2a* mRNAs in hippocampal neurons, we generated the *Camk2a* stem loop knock-in mouse model, referred to as *Camk2a*-*MS2/PP7* KI, where an array of optimized 24 alternating MS2 and PP7 stem loops (i.e. 24x MS2/PP7) were knocked into the end of 3’untranslated regions (UTRs) in the *Camk2a* alleles. Exogenous expression of green fluorescent protein (GFP) or HaloTag-fused coat proteins that bind to the stem loops enabled direct visualization of endogenous *Camk2a* mRNA granules in living hippocampal neurons cultured from the *Camk2a*-*MS2/PP7* KI mice (Figure1A, Supplementary Movie1). *Camk2a* mRNA granules under the basal condition displayed previously observed mRNA movements, including anterograde, retrograde and corralled movements, suggesting that the mRNAs were transported by microtubule-based mechanisms (*26, 27*). Single molecule *in situ* fluorescence hybridization (smFISH), probing for the endogenous *Camk2a* mRNAs and stem loop-tagged mRNAs in hippocampal neurons of homozygous *Camk2a*-*MS2/PP7* KI mice, showed that the majority of the stem loop signal was co-localized with the *Camk2a* mRNA signal within 250 nm distance, and the frequencies of co-localization among randomly sampled dendritic segments were positively correlated (r=0.69), indicating that the integrity of the mRNAs containing intact stem loops was maintained (Supplementary Figure 1A-C) (*28, 29*). smFISH showed that untagged and stem loop containing *Camk2a* mRNAs were equally present in heterozygous hippocampal neurons and that neuronal activation by withdrawal of the voltage-gated sodium channel blocker tetrodotoxin (TTX) increased mRNAs abundance in the proximal dendrites for both tagged and untagged mRNAs (Supplementary Figure 1E). This result suggested that addition of exogenous stem loops to the *Camk2a* 3’UTR did not alter stability, transport or localization of the endogenous mRNAs (Supplementary Figure 1D-E). Therefore, this mouse model can serve to understand the activity-dependent localization dynamics of an mRNA involved in synaptic plasticity.

To determine the activity-dependent localization of endogenous *Camk2a* mRNAs with high spatial and temporal resolution, distal dendritic regions were locally stimulated using photolytic uncaging of glutamate that binds to NMDARs on the postsynaptic dendritic spines. Uncaging of glutamate induced rapid and robust localization of the *Camk2a* mRNA granules to ∼80% of uncaged extradendritic spots with an average arrival time of ∼85 seconds (Figure 1B-D). Uncaging-induced localization patterns indicated that *Camk2a* mRNAs were situated perpendicular to the axes of dendritic shafts. The corralled movements of the mRNAs suggested that they were restricted in confined spaces 0.5-1 micrometers away from the dendritic shaft, possibly in dendritic spines (Figure 1C). sLTP is a temporally ordered process, starting with the activation of postsynaptic CaMKII by Ca^2+^ and calmodulin (CaM) to structural changes, involving actin polymerization, that follow in minutes and hours (*30, 31*). The timing of uncaging-induced mRNA localization was in line with those of the maintenance-related processes (Figure 1D and Supplementary Movie 1) (*8, 32, 33*). The recruitment of *Camk2a* mRNAs to the uncaged spots was robust; ∼80% of uncaged spots recruited discrete mRNAs granules (Figure 1D). Together, live imaging of endogenous *Camk2a* mRNAs in mouse hippocampal neurons revealed dynamic activity-dependent mRNA localization to potential postsynaptic dendritic spines.

**Figure 1.**
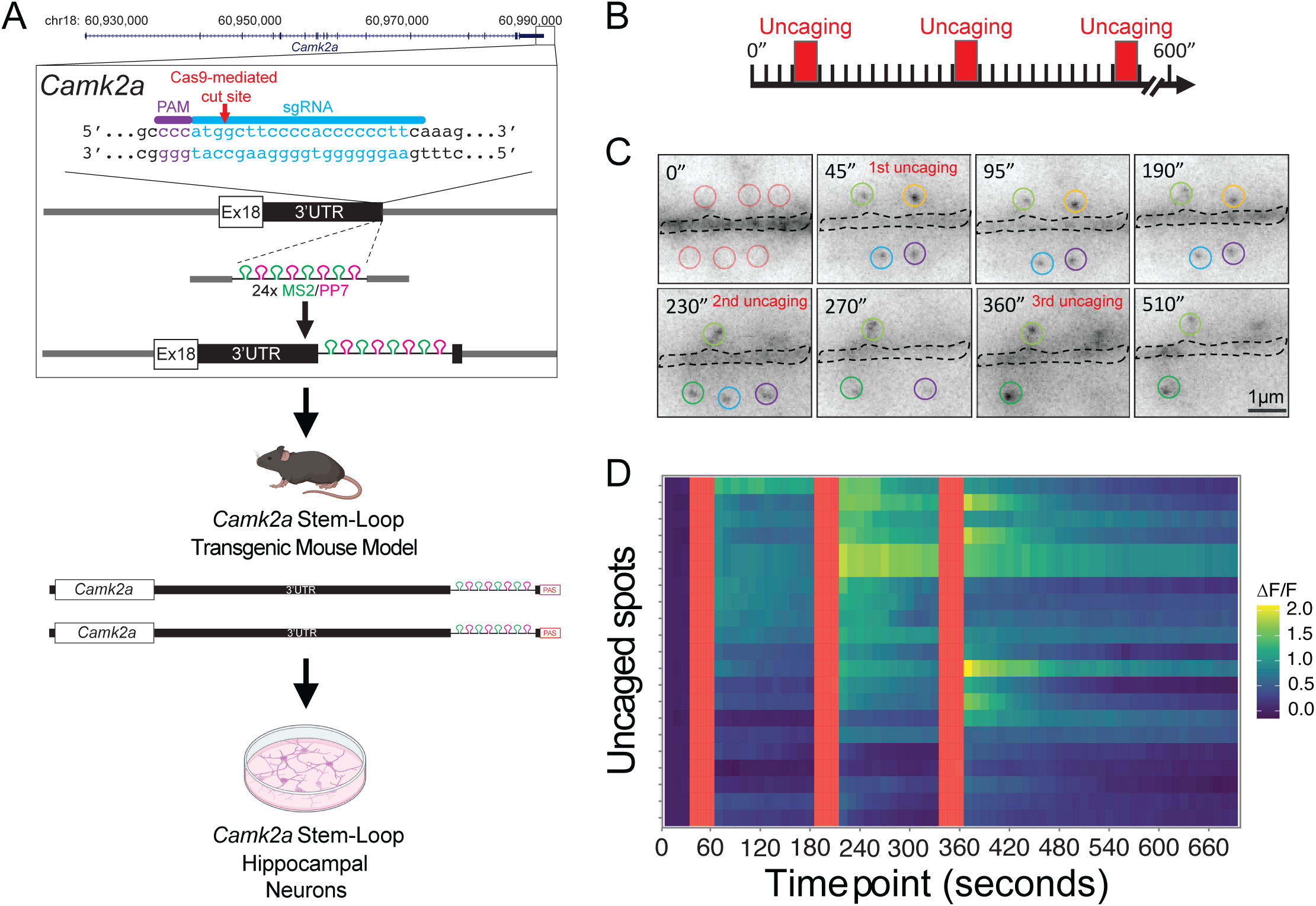
Generation of *Camk2a* stem-loop knock-in mouse and localization of *Camk2a* mRNA at the postsynaptic dendritic spines. (**A**) CRISPR/Cas9-mediated knock-in strategy for generating the *Camk2a* stem-loop transgenic mouse that is maintained at homozygosity and implementation of hippocampal neuron culture. (**B**) Low-frequency glutamate uncaging protocol in Mg^2+^-free and Tetrodotoxin (TTX)-supplemented medium with 3 trains of 10 pulses at 0.5Hz were applied to individual dendritic spines. (**C**) Representative time series illustrates the *Camk2a* mRNA signals in six uncaged spots (demarcated by colored circles) at marked time points post-glutamate uncaging. (**D**) Quantification shows the fluorescence intensity changes (Δ**F**) over the initial background (**F**) of mRNA signals in individual uncaged spots (Each row displays the intensity changes, n=21).

### Activity-dependent localization of the *Camk2a* mRNAs to the postsynaptic density in dendritic spine heads

The plausible location of glutamate-induced *Camk2a* mRNA localization was inside postsynaptic dendritic spines based on the geometry and confined movements of detected mRNAs (Supplementary Movie 1, 2). To establish this, we examined the degree of association between the postsynaptic scaffolding protein PSD95 (proxy for the PSD) and dendritic *Camk2a* mRNAs at various time points after chemical long-term plasticity (cLTP): unstimulated control (basal), initial phase (5 minutes post stimulation), and maintenance phase (45 minutes post stimulation), in the presence or absence of a selective NMDAR antagonist D-APV (Figure 2A-B). Simultaneous smFISH and immunofluorescence staining (smFISH-IF) allowed co-detection of *Camk2* mRNAs along with PSD95 and dendritic protein marker MAP2. The degree of association was computed as the frequency of the PSD signals that associated with *Camk2a* mRNAs within the 2D radial distance of 500 nm in distal dendritic segments. Despite basal association of *Camk2a* mRNA with PSD in unstimulated neurons (12.92% ± 1.39%, Mean ± S.E.M.), significant increases to 20.52% ± 1.52% (*p*-value = 2.00E-02) immediately after stimulation (5 min post-cLTP) and to 24.03% ± 1.72% (*p*-value = 5.30E-04) at later time point (45 min post-cLTP) were observed, indicating that NMDAR activation led to enrichment of *Camk2a* mRNA to the PSD of dendritic spines (Figure 2A-B). Notably, the enrichment of *Camk2a* to the PSD was significantly reduced in the presence of D-APV (9.60% ± 1.00% and 9.91% ± 1.02% at 5 min and 45 min post-cLTP, respectively), indicating that mRNA localization was driven by the activation of NMDARs. Cross pair correlation functions computed the probabilities to find *Camk2a* mRNA and the PSD signals at varying inter-signal distances further showed that while there was a general enrichment of *Camk2a* mRNA to the PSD even under basal conditions (possibly due to network activity in the cultures), the highest probability was found in stimulated neurons (Supplementary Figure 2). The average size of the *Camk2a-*associated PSD in stimulated neurons (0.41μm^2^ ± 0.06μm^2^; *p*-value *=* 1.16E-11) was approximately twice as large as those that were not associated with the mRNAs (0.28μm^2^ ± 0.01μm^2^), illustrating that cLTP-induced *Camk2a* localization preferentially occurred to more mature dendritic spines undergoing structural plasticity (Figure 2C). Importantly, *Camk2* mRNA’s propensity to localize to the larger PSD was abrogated in the presence of D-APV (0.29μm^2^ ± 0.03μm^2^ vs. 0.25μm^2^ ± 0.01μm^2^; *Camk2a-*associated vs. *Camk2a-*unassociated). Next, to investigate whether *Camk2a* mRNA localization to the PSD leads to concomitant structural changes in the spines, time lapse imaging of *Camk2a* mRNAs and spine morphology (ABD-mCherry) was performed 30 min after cLTP. Indeed, *Camk2a* mRNAs enriched spines underwent significant enlargement in size with higher actin polymerization over time compared to the spines without mRNAs (Figure 2D-F). These results together with local glutamate-induced localization (Figure 1B-D) suggested that *Camk2a* mRNAs localized to dendritic spines near the PSD via activation of NMDARs, and this activity-dependent spine localization was positively correlated with the size of PSD, linking mRNA localization to the enhancement of synaptic strength of individual dendritic spines.

**Figure 2.**
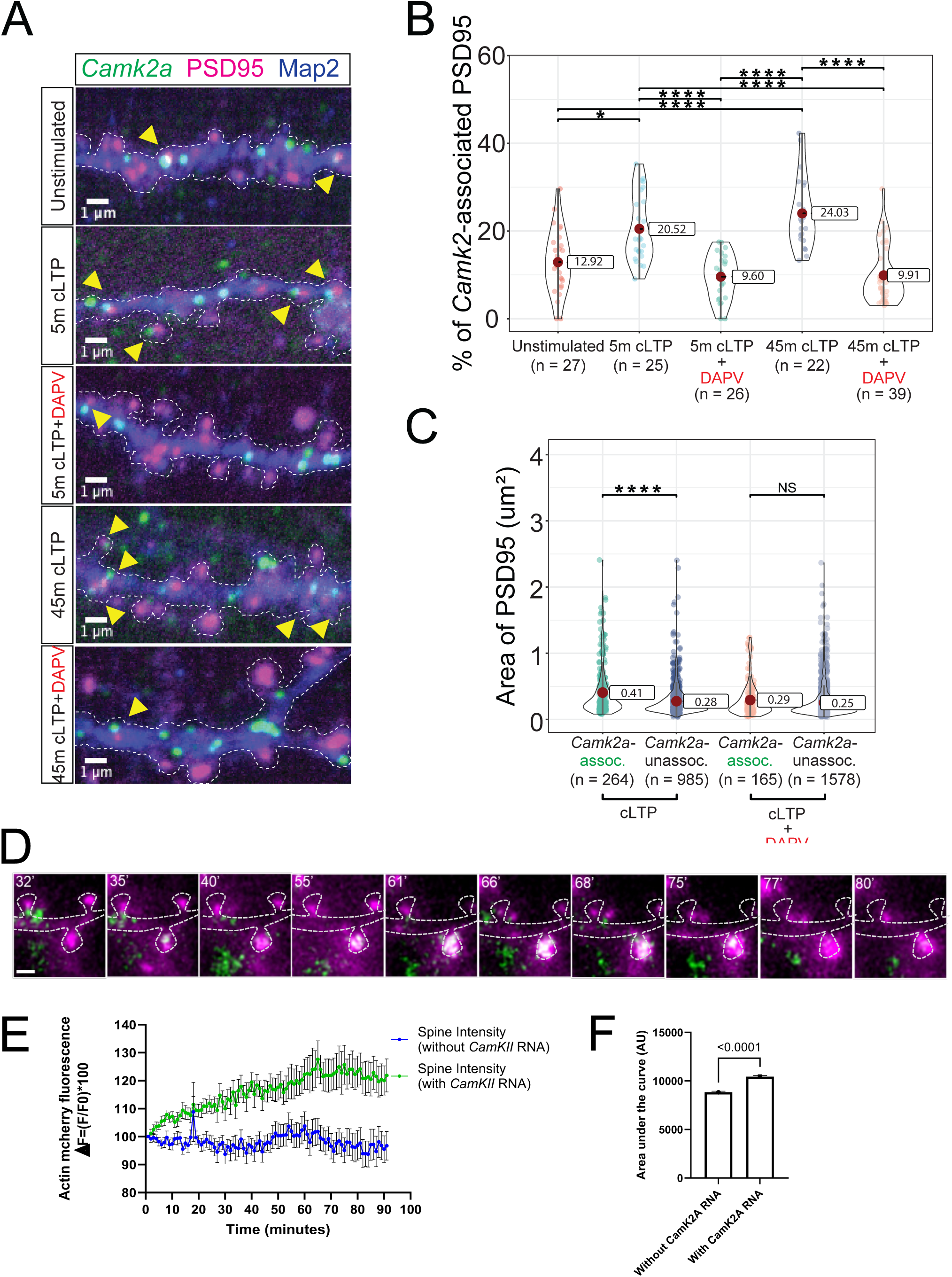
Activity-dependent localization of *Camk2a* mRNAs to the postsynaptic density PSD95 in mouse hippocampal neurons. (**A**) Representative images display localization of *Camk2a* mRNAs (green) to the postsynaptic PSD95 (magenta) and in dendrites (blue) of unstimulated (basal), stimulated (Glycine 200μM + Picrotoxin 100μM), and stimulated neurons in the presence of a selective NMDAR antagonist D-APV (50nM). (**B**) Quantification shows the mean frequencies of the *Camk2a* localization to PSD95 in corresponding conditions: unstimulated, 5 min post-cLTP, 5 min post-cLTP in the presence of D-APV, 45 min post-cLTP and 45 min post-cLTP in the presence of D-APV. (the indicated number of datapoints refers to the number of analyzed cells). Statistical significances are estimated by One-way ANOVA and indicated by horizontal bars. *p*-value annotation: * ≤0.05, **≤0.01, ***≤0.001, ****≤0.0001 (**C**) Quantification shows the estimated sizes of the *Camk2a*-associated and unassociated PSDs among unstimulated, stimulated by cLTP and stimulated by cLTP in the presence of D-APV (the indicated number of datapoints refers to the number of analyzed PSD95 signals). Statistical significance is estimated by paired t-test. *p*-value annotation: * ≤0.05, **≤0.01, ***≤0.001, ****≤0.0001 (**D**) Time-lapse images of *Camk2a* mRNAs localizing to spine heads in specific spines associated with spine enlargement. (green = *Camk2a* mRNAs, magenta = Actin-mCherry). Timestamps indicate post cLTP stimulation time. Scale bar = 1 micron. (**E**) Quantification of spine actin intensity change in spines containing *Camk2a* mRNAs and the neighboring ones without RNA localization. (**F**) Area under the curve from (E). *p*-value < 1.0E-04, Mann Whitney test.

### *Cis*-acting regulatory elements CPEs in 3’UTR of *Camk2a* confer activity-dependent spine localization of the mRNAs

To identify the *cis*-acting regulatory elements in the *Camk2a* mRNA that drive spine localization, we focused on the CPE regulatory sequences on the 3’UTR, which are recognized by cognate RNA binding proteins, the cytoplasmic polyadenylation element-binding proteins (CPEBs). Importantly, CPEBs are known to regulate activity-dependent polyadenylation and protein synthesis of synaptic mRNAs (*16, 34, 35*). We first tested the requirement of CPEs for spine localization of *Camk2a* mRNAs by using the morpholino antisense oligonucleotides (ASOs) targeting CPE elements. CPE-targeting ASOs (i.e. CPE ASOs) consisted of two ASOs that were designed to stably bind to tandem CPE elements in the 3’UTR of *Camk2a* mRNA to obstruct CPE-mediated functions (Figure 3A). Non-targeting ASO (i.e. Control ASO) was used as a control. To minimize indirect effects of ASOs, we transiently introduced ASOs to *Camk2a-MS2/PP7* KI hippocampal neurons 2 hours before the experiment. We then assessed the activity-dependent spine-localization frequencies of *Camk2a* mRNA signals by quantifying mRNAs localized within 500 nm radial distance of GFP-CaMKIIα-filled spine heads (Figure 3B and Supplementary Movie 3). While the baseline spine-localization frequencies between Control ASO and CPE ASOs-treated groups were comparable (13.62% ± 1.39% vs. 10.5% ± 1.57%), the corresponding frequencies in stimulated neurons at 45 minutes post-stimulation significantly differed (26.11% ± 1.74% vs. 15.24% ± 1.78%; Control ASO vs. CPE ASOs; *p*-value *=* 8.13E-04), in that the pretreatment of neurons with CPE ASOs dampened the spine-localization frequency (Figure 3B-C). This result indicated that CPEs in the 3’UTR of *Camk2a* were necessary for activity-dependent localization of the mRNAs to dendritic spine heads.

**Figure 3.**
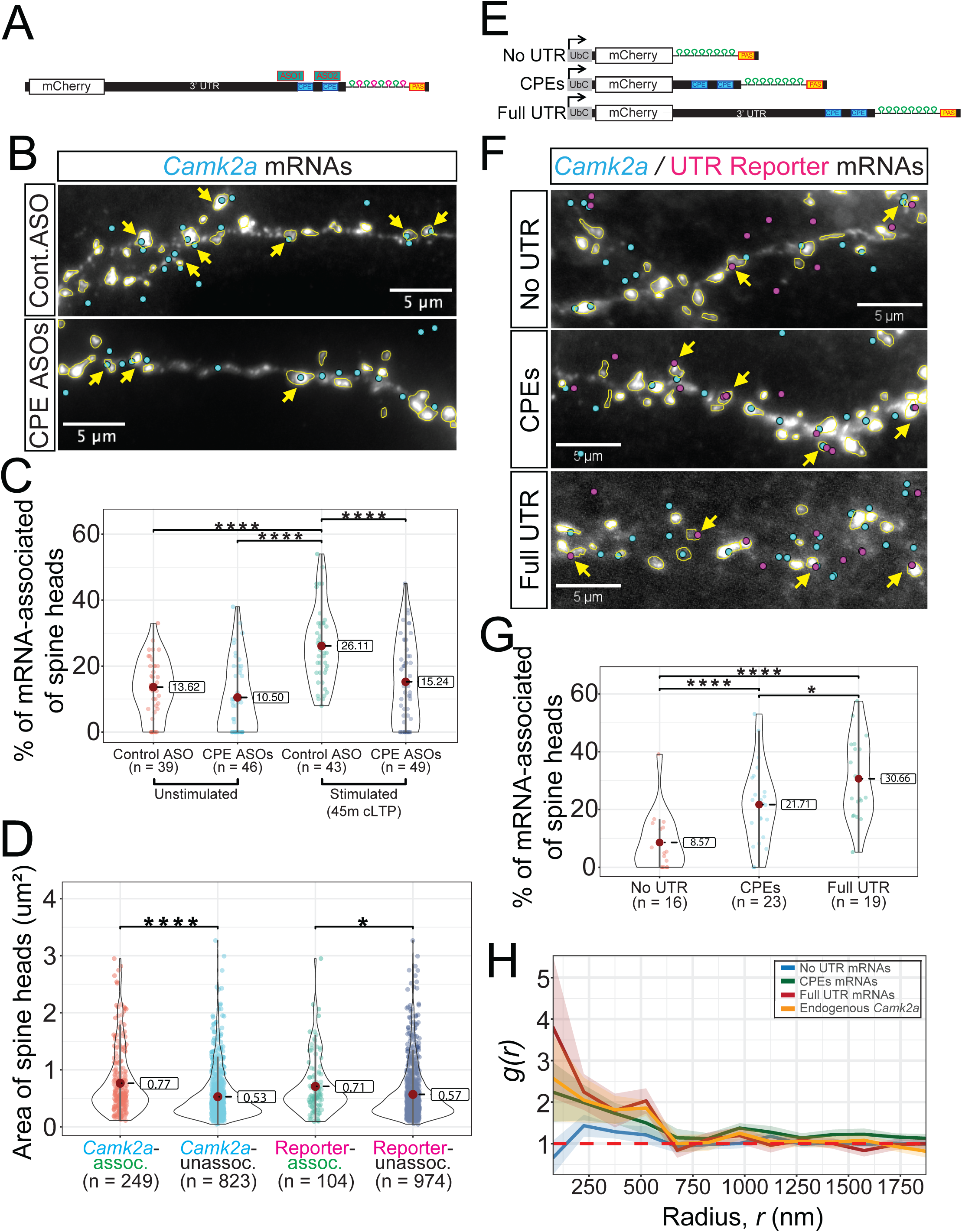
*Cis*-acting CPEs in 3’UTR of *Camk2a* confer activity-dependent spine-localization of the mRNAs. (**A**) Schematic illustrates the design of morpholino antisense oligos (ASOs) against *cis*-acting CPEs. (**B**) Representative images show localization of the *Camk2a* mRNAs (cyan) to the GFP-CaMKIIα-filled dendritic spines (demarcated by yellow outlines) in stimulated neurons at 45 minutes post-cLTP, which are pretreated with either non-targeting control ASO or CPE-targeting ASOs. Yellow arrows indicate the spine-localized *Camk2a* mRNAs (**C**) Quantification shows the mean frequencies of the *Camk2a* localization to spine heads in corresponding conditions (datapoints refers to individual neurons). Statistical significances are estimated by one-way ANOVA and indicated by horizontal bars. *p*-value annotation: * ≤ 0.05, ** ≤ 0.01, *** ≤ 0.001, **** ≤ 0.0001 (**D**) Pairwise comparisons of the estimated sizes of the endogenous *Camk2a*-associated vs. mRNA-unassociated spine heads and 3’UTR reporter-associated vs. reporter-unassociated spine heads at 45 minutes post-cLTP (n = number of analyzed spine heads). Statistical significance is estimated by paired t-test. *p*-value annotation: * ≤ 0.05, ** ≤ 0.01, *** ≤ 0.001, **** ≤ 0.0001 (**E**) Schematic illustrates the design of reporter mRNA constructs that harbor no UTR (No UTR), 2 CPEs (CPEs) and *Camk2a*’s 3’UTR (Full UTR) in conjunction with MS2 stem loops. (**F**) Representative images show localization of the endogenous *Camk2a* mRNAs (cyan) or the exogenous 3’UTR reporter mRNAs (magenta) to the GFP-CaMKIIα-filled spine heads (demarcated by yellow outlines) in stimulated neurons at 45 minutes post-cLTP. Yellow arrows indicate the spine-localized 3’UTR reporter mRNAs (**G**) Quantification shows the adjusted frequencies of spine localization of the 3’UTR reporter mRNAs in corresponding groups (n = number of analyzed cells). Statistical significances are estimated by one-way ANOVA and indicated by horizontal bars. *p*-value annotation: * ≤0.05, **≤0.01, ***≤0.001, ****≤0.0001 (**H**) Quantification shows cross correlation function (*g(r)*) of 3’UTR reporter mRNAs localized to centroids of spine heads, represented as a function of inter-signal distances between the pairs (radius, *r*) at 45 minutes post-cLTP. *g(r)* >1 indicates a significant probability to find two signals at a given radius.

Next, we tested whether CPEs were sufficient to drive spine mRNA localization by examining activity-dependent spine localization of 3’UTR reporter mRNAs that harbored CPE sequences in their 3’UTR (Figure 3E). We devised three reporter mRNA constructs: no UTR reporter where the 3’UTR was omitted, a CPE reporter that had a minimal CPE-containing segment of the *Camk2a* 3’UTR, and a full UTR reporter equipped with the entire *Camk2a* 3’UTR. These 3’UTR reporter constructs were transduced into mouse hippocampal neurons, and the spine-localization frequencies of reporter mRNAs in relation to GFP-CaMKIIα-filled spine heads at 45 minutes post-cLTP stimulation were calculated (Figure 3F-G). Indeed, reporter mRNAs containing CPEs localized to dendritic spines (21.71% ± 2.85% and 30.66% ±3.22%; CPE and full UTR; *p*-value = 8.68E-03 and *p*-value = 6.00E-05, respectively) with significantly higher frequencies than no UTR control reporter (8.57% ± 2.58%), indicating that CPEs in the 3’UTR were sufficient to drive spine localization of the respective mRNA granules (Figure 3G). Consistent with previous findings, the average size of endogenous *Camk2a*-associated spine heads (0.77μm^2^ ± 0.04μm^2^; *p*-value = 9.12E-12) was larger than the average size of *Camk2a*-unassociated spine heads (0.53μm^2^ ± 0.01μm^2^) (Figure 3D). Of note, the 3’UTR reporter mRNA were associated with larger spines (0.71μm^2^ ± 0.05μm^2^; *p*-value = 2.00E-02, compared to 0.57μm^2^ ± 0.01μm^2^ for reporter mRNA-unassociated spine heads) (Fig 3G), suggesting that the parameters such as the maturity or the potency of spines are intrinsic factors that promote or result from mRNA localization. Lastly, cross pair correlation revealed higher probability of finding mRNAs that contained CPEs (i.e. CPEs, Full UTR and endogenous *Camk2a*) within 500 nm radial distance of the spine heads in comparison with mRNAs lacking CPEs (No UTR), reiterating that CPEs sufficiently drove spine localization of mRNAs (Figure 3H). Together, these findings provided evidence that the CPEs were necessary and sufficient for activity-dependent localization of *Camk2a* mRNAs to the spine heads.

### Characterization of activity-dependent local protein synthesis of *Camk2a* mRNA in dendritic spines

The implication of activity-dependent and CPE-mediated localization of *Camk2a* mRNA to dendritic spine heads is on-site protein synthesis. Neuronal activity induces the spatiotemporal compartmentalization of local environments, therefore, how proteins are made in distinct dendritic compartments is of great importance (*36*). Polyribosomes are found in the spine heads, however, real-time detection of mRNA translation in spines has not been possible before (*37, 38*). We therefore sought to directly visualize translating *Camk2a* mRNAs by monitoring the nascent CaMKIIα peptide chains as they are being synthesized from the mRNA molecules in response to activation of NMDARs. To capture nascent chains and the mRNA molecules simultaneously, we knocked in an array of SunTag epitopes to the 5’ end of the *Camk2a* coding sequence in hippocampal neurons that were cultured from homozygous *Camk2a*-*MS2/PP7* KI mice (Supplementary Figure 4A). A successful Cas9-mediated genome editing yielded the transgenic allele that consisted of SunTag epitopes, *Camk2a* coding sequence and the stem-loop within a single mRNA.

Using this system, we found fractions of co-localizing mRNA and SunTag signals in both dendrites and dendritic spines 15 minutes after cLTP (Figure 4A-C). Treatment with the protein synthesis inhibitor Cycloheximide (CHX) showed a sharp decline in SunTag signal without affecting the mRNA signal, suggesting that SunTag signal is a robust readout of ongoing CaMKIIα translation events (Figure 4A-B). Translating mRNAs were detected in both dendritic shafts and inside the spine heads but the probability of translation was significantly higher inside the spine heads than in the shafts (73.6 ± 6.3 % in spines vs 15.5 ± 5.6 % in shafts, *p*-value = 8.00E-04, Figure 4C-D). Moreover, the translation continued inside the spines for a longer duration compared to that in the dendritic shafts (Figure 4E), indicating that the efficiency of local translation depended on the specific compartment where mRNA was localized. A considerable fraction (∼60%) of mRNA and SunTag signals remained at the end of time-lapse imaging, indicating the persistent presence of both *Camk2a* mRNA and SunTag peptides in each sub-compartment despite some photobleaching (Supplementary Figure 4B-E). The lifetime estimation revealed the extents to which SunTag peptides in dendrites and dendritic spines could remain locally over several hours (Supplementary Figure 4D-E). In comparison with *Camk2a* mRNAs in dendrites, those in dendritic spines appeared to decay slower, thus they were more persistent (Supplementary Figure 4B-C). These findings highlighted the dendritic spines as the sites of neuronal activity-induced local protein synthesis, where *Camk2a* mRNA localization was spatially coupled to translation and this on-site translation could last for extended periods, consistent with their roles in the long-term plasticity.

**Figure 4.**
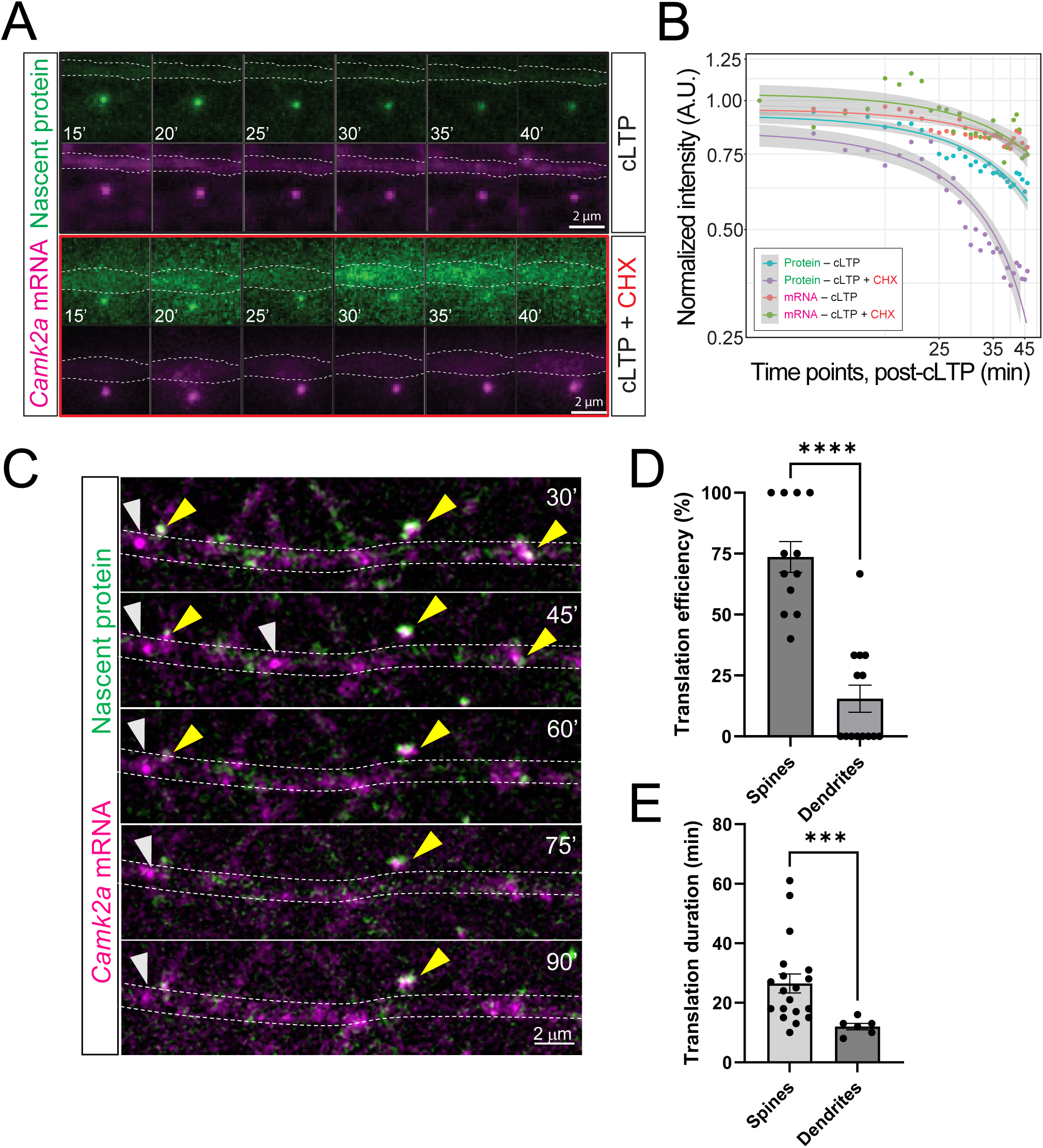
Activity-induced local protein synthesis of *Camk2a* mRNA in dendrites and dendritic spines by simultaneously detecting the nascent SunTag-CaMKIIα peptides and *Camk2a* mRNA. (**A**) Representative time series illustrates the SunTag nascent protein (green) and *Camk2a* mRNA (magenta) at dendritic spines 45 minutes post cLTP in the absence (upper panels) and the presence (lower panels) of cycloheximide (CHX). (**B**) Quantification of normalized fluorescence signal intensities show the extended co-localization of the nascent SunTag proteins and corresponding mRNA signals over 45 minute-period and dissipation of the SunTag signals in the presence of CHX (Each data point displays the means of normalized signal intensities, n=8 events from 3 independent experiments). (**C**) Translation of *Camk2a* mRNAs in dendrites following cLTP stimulation (yellow arrows indicate translating mRNAs, white arrows indicate non-translating *Camk2a* mRNAs). (**D**) Translation efficiency measured by fraction of mRNAs associated with SunTag signals, n = 12 dendrites from 3 independent experiments. *p-*value = 8.0E-04, Mann Whitney test. (**E**) Quantification of translation duration. n = 19 events for spines, n=6 events for dendritic shafts. *p-*value < 1.0E-04, Mann Whitney test.

## Discussion

In this work, we report a novel activity-dependent local regulation of *Camk2a* mRNA in mouse hippocampal neurons, in which the mRNA rapidly localized to stimulated dendritic spines, positioned near the PSD and underwent on-site translation preferentially inside the spines. By generating a knock-in mouse where all the endogenous *Camk2a* mRNAs were tagged with stem loops and fluorescently labeled, we were able to obtain high resolution insight into *Camk2a* mRNA dynamics and uncover a hereto unknown localization that explains how local CaMKIIα protein can be maintained in specific spines undergoing plasticity for extended periods of time. The timescale of the localization was rapid (∼85 seconds) for *Camk2a* mRNA to localize to ∼80% of the uncaged regions. The rapid arrival of *Camk2a* to stimulated spines suggests that the mRNA localization was an immediate response to the initial Ca^2+^ signal following NMDAR activation. Once localized, the mRNAs resided in the spines for up to 60 minutes. The time window of this persistent localization of *Camk2a* mRNA overlapped with the maintenance phase of long-lasting synaptic plasticity (*2*). The spatial scale of this localization of *Camk2a* was unequivocally specific to dendritic spines as the mRNAs were enriched within 500 nm of the PSD, suggesting that the localization was singularly targeted to stimulated spines. Importantly, this spatial specificity was lost in the presence of the competitive NMDAR antagonist D-APV, indicating that the process relied on activity of the NDMAR – hence was activity-dependent. In addition, *Camk2a* mRNA tended to be associated with larger PSD, where more α-amino-3-hydroxy-5-methyl-4-isoxazolepropionic acid receptors (AMPARs) can be located, suggesting that the spine maturity and the potentiation capacity may be the intrinsic determining factors in this process (*39, 40*). The real time measurements of spine size underscored that *Camk2a* mRNA localization could drive significant actin polymerization-mediated enlargement in those spines compared to neighboring ones lacking persistent RNA localization. This highlights the uniqueness of the activity-dependent spine localization of *Camk2a* mRNA and its role in promoting structural plasticity in specific spines.

In neurons, local regulation of synaptic mRNAs often involves two spatiotemporally coordinated processes: (1) localization of the mRNAs to the sites near stimulated synapses and (2) ensuing local protein synthesis. A collective goal of these local regulatory mechanisms is thought to situate stable sources of synaptic proteins near the synapses of action (*26, 27, 41–43*). For example, patrolling β-actin mRNAs become docked at the bases of stimulated spines and undergo local protein synthesis whereas Arc mRNAs, replenished by cyclic activity-dependent transcription, make transitory, but frequent stops in the vicinity of the stimulation sites to assemble Arc protein hubs (*21, 23*). To determine the behavior of *Camk2a* mRNAs, we devised a *Camk2a*-*MS2/PP7* KI mouse model, which harbors an array of optimized MS2/PP7 hybrid stem loops in the 3’UTRs of the endogenous *Camk2a* alleles. This animal model has allowed us to probe detailed spatiotemporal distribution patterns of the endogenous *Camk2a* mRNAs in relation to the postsynaptic dendritic spines following plasticity induction and maintenance. By implementing a recently developed CRISPR/Cas9-mediated knock-in approach and the real-time nascent SunTag epitope tagging method on this mouse model, we resolved not only individual mRNAs but also newly synthesized protein polypeptides as discrete fluorescent signals in spatially restricted subcellular compartments such as postsynaptic dendritic spines (*44, 45*). These technological advancements provided us with the means to investigate how spine-recruited *Camk2a* localization was spatiotemporally choreographed with local protein synthesis.

Previous studies have shown that a genetic disruption of the 3’UTR of *Camk2a* substantially depletes the dendritic pool of the mRNA in mouse hippocampus and neocortex, which in turn leads to more than 50% reduction in the overall level of CaMKIIα protein in the respective brain homogenates, thus underscoring a critical role of the 3’UTR and dendritic local protein synthesis in protein homeostasis (*20, 46*). We noted that the polyadenylation-related *cis*-acting regulatory elements CPEs in the 3’UTR of *Camk2a* mediates activity-dependent local protein synthesis, the mechanism of which is attributable to the polyadenylation implemented by cognate RNA-binding protein CPEB (*16, 34*). Importantly, *Camk2a* and CPEB seemingly converged on dendritic spines via CPEs (*16, 34, 35*). Indeed, the spine-localization of *Camk2a* was mediated by CPEs as targeted obstructions of the elements using the stable morpholino ASOs disrupted the spatial enrichment of mRNA in dendritic spine heads. Conversely, when located in the 3’UTR, these elements promoted activity-induced spine localization of the reporter mRNAs, thus proving necessity and sufficiency of CPEs. Of note, while interaction between 3’ UTR of *Camk2a* mRNA and CPEB proteins (i.e. CPEB1) via CPEs has been demonstrated *in vitro*, it has yet to be directly visualized in neurons (*16*). In addition, it remains unknown whether other three neuronal CPEB protein family members (e.g. CPEB2-4) can interact with *Camk2a*, thereby participating in spine localization (*47*). Our analysis showed that when CPEs were present on their own as a minimal segment, the efficiency of CPE-mediated spine localization was suboptimal, suggesting that other *cis*-acting regulatory sequences elsewhere in the 3’UTR may still be required to obtain an optimal spine-localization efficiency (see Figure 3G).

The major implications of our finding of CPE-mediated spine localization of *Camk2a* mRNA included activity-induced local protein synthesis at dendritic spines. In fact, we demonstrated that the extended presence of *Camk2a* mRNA at stimulated dendritic spines was accompanied by a robust and persistent on-site protein synthesis throughout mRNA’s stay. Thus, *Camk2a* localization and corresponding protein synthesis were tightly coupled in time and space, one of the first demonstrations of efficient coupling mechanism for a dendritically localized mRNA. This explicit spine local regulation of *Camk2a* mRNA may represent a plausible outcome in establishment of synapse-specific pool of CaMKIIα. Persisting on-site protein synthesis is a potential mechanism that could give rise to a steady elevation in local protein concentration in a spatially restricted dendritic spine near the PSD. High concentration of proteins would be expected to favor incorporation of newly synthesized CaMKIIα subunits into the pre-existing holoenzymes by shifting the equilibrium between association and dissociation in a spatially contained manner. In fact, it has been demonstrated *in vitro* that activation of CaMKIIα increases the intermolecular subunit exchange among holoenzymes contained in the same space (*48, 49*). A persistent subunit incorporation would lead to the perpetuation of synaptic transmission at a given synapse, thereby providing a long-term basis for maintaining synaptic activity in the wake of transient Ca^2+^ signal (*50*). In this way, activity-dependent mRNA localization and local translation for the plasticity gene *Camk2a* is a mechanism for long term protein maintenance at synapses required for LTP.

## Materials and Methods

### Generation of *Camk2a* stem-loop knock-in mouse

All animal care and experimental procedure were implemented in accordance with the protocols approved by the Institutional Animal Care and Use Committee (IACUC) at Albert Einstein College of Medicine and Janelia Research Campus of Howard Hughes Medical Institute (HHMI). The *Camk2a*-*MS2/PP7* knock-in (KI) mouse model was generated by integration of an array of 24 optimized repeats of alternating MS2 and PP7 stem loops (24x MS2/PP7) to the end of 3’untranslated region (3’UTR) of the endogenous *Camk2a* allele. This newly designed sequence contained randomized linker sequence between the MS2 and PP7 loops to prevent recombination events. Using CRISPR/Cas9 technology and a guide RNA (gRNA) targeting the 3’ end region of mouse *Camk2a*’s 3’UTR (gRNA sequence: 5’-AAGGGGGGTGGGGAAGCCAT-3’), *Camk2a*-*MS2/PP7* KI allele was created in 129S6 × C57BL/6 F1 hybrid mouse embryonic stem (ES) cell line. BL/6 blastocysts injected with targeted ES cells, harboring the KI allele, were then transferred into pseudopregnant female mice and generated chimeras. The resulting progenies from mating between the chimeras and C57BL/6 wild type mice were screened for the integration of the endogenous *Camk2a*-*MS2/PP7* KI allele in the correct orientation by PCR and DNA sequencing. Finally, mice with successful germline transmission of the allele were backcrossed to C57BL/6 wild type mice to establish the *Camk2a*-stem loop KI mouse lines. Mice that were backcrossed into C57BL/6 background at least 10 times or more were used for all experiments.

### Preparation of mouse hippocampal neurons

Dissociated hippocampal neuron culture was prepared as previously described (*1, 2*). Briefly, hippocampi collected from 4-6 brains of homozygous, heterozygous *Camk2a*-*MS2/PP7* KI or C57BL/6 wild type mouse pups at postnatal day 1-2 (P1-2) were pooled and digested with 0.25% Trypsin (Gibco) in 1x HBSS (Thermo Fisher) buffer, supplemented with 10mM HEPES (Thermo Fisher). Dissociated hippocampal neurons were seeded onto the poly-D-lysine (Sigma-Aldrich) coated surface of glass bottom cell culture dishes (MatTek) and incubated in neurobasal A (Thermo Fisher)-based neural growth medium (NGM), supplemented with 10% fetal bovine serum (R&D systems), L-glutamine alternative GlutaMAX (Thermo Fisher) and antimicrobial Primocin (InvivoGen), at 37°C and 5% CO_2_ for 2 hours. Following the initial incubation, hippocampal neurons were subsequently cultured in NGM, supplemented with 1% fetal bovine serum, B27 (Thermo Fisher), GlutaMAX and Primocin for up to 4 weeks (i.e. days in vitro 28, or DIV 28). All experiments were performed, using cultured hippocampal neurons at DIV 21-28.

### Visualization of *Camk2a* mRNA, SunTag-CaMKIIα nascent protein and MS2 stem loop-tagged reporter mRNAs in living hippocampal neurons with lentiviral systems

To visualize the stem loop-tagged *Camk2a* mRNA molecules alone in living hippocampal neurons, following lentiviral vectors, housing coding sequences of fluorescently labeled coat proteins, were transduced into cultured neurons as previously described: (1) the synonymized versions of tandem PP7 coat proteins and two green fluorescent proteins (stdPCP-stdGFPs), (2) the synonymized tandem MS2 coat proteins and two stdGFPs (stdMCP-stdGFPs) and (3) stdMCP and a HaloTag (stdMCP-HaloTag) along with Jenelia fluor dyes(*1–3*). To generate lentiviral particles, lentiviral plasmids, compatible lentiviral packaging plasmids (pMDLg/pRRE, pRSV-Rev) and envelope plasmids (pMD2.VSVG) were transfected into HEK293T cells using jetPRIME transfection reagent (PolyPlus). The resulting supernatant medium, containing lentiviral particles, was collected and concentrated using Lenti-X concentrator (Takara). Expression levels of all lentiviral vectors, expressing fluorescently tagged proteins and single chain antibodies, were optimized by an adjustment of viral titers.

For simultaneous visualization of *Camk2a* mRNA and SunTag-CaMKIIα nascent protein molecules, first we employed CRISPR/Cas9-based endogenous knock-in approach to integrate an array of 23 repeats of SunTag (i.e. GCN4) epitopes in frame with *Camk2a* coding sequence at the 5’ end at the endogenous level. To this end, recently developed pORANGE CRISPR/Cas9 system (*4*), which was originally designed to fuse fluorescent proteins (e.g. EGFP and mEos3.2) to the endogenous CaMKIIα at the N-terminus, was modified so that the original EGFP coding insert sequence was replaced with 23 repeats of SunTag epitopes. Briefly, 2ug of SunTag-containing pORANGE CRISPR/Cas9 knock-in plasmid DNA was transiently transfected into hippocampal neurons at DIV5, using Lipofectamine 2000 reagent (Thermo Fisher) at 1ul/ml of media. To visualize both *Camk2a* mRNA and SunTag-CaMKIIα molecules in the same transduced neurons, a bicistronic lentiviral vector, in which coding sequences of stdMCP-HaloTag and single chain antibody fragment fused with the superfolder GFP (scFv-sfGFP) were partitioned by internal ribosome entry site (IRES) of the encephalomyocarditis virus (EMCV), was transduced into hippocampal neurons. Successfully CRISPR/Cas9 knock-in of the SunTag array and transduction of stdMCP-HaloTag; scFv-sfGFP bicistronic vector was confirmed by the concurrent detection of mRNA and SunTag signals.

For an assessment of the role of *Camk2a*’s 3’UTR CPEs in activity-dependent spine-localization, iterations of lentiviral mRNA reporter plasmids were constructed as follows: (1) No UTR reporter had coding sequence of a placeholder mCherry followed by 24 repeats of MS2 stem loop (i.e. MS2V5) only, (2) CPEs reporter had mCherry coding sequence followed by MS2V5 and the last 436 bp-long segment of rat *Camk2a*’s 3’UTR, encompassing two previously characterized CPEs, and (3) Full UTR reporter had coding sequence of mCherry followed by the first 2920 bp-long segment of rat *Camk2a*’s 3’UTR, MS2V5 and the last 436bp-long CPE-containing segment. Successful transduction of these mRNA reporter vectors in hippocampal neurons was validated by the detection of MS2V5-tagged mRNA signals in the presence of exogenously expressed stdMCP-stdGFP in living neurons and specific smFISH probes against MS2V5 stem loops in fixed neurons.

### Live imaging of stimulated hippocampal neurons

Photolytic uncaging of MNI-caged-L-glutamate (Tocris) was employed to stimulate NMDARs on dendritic spines on small segments of dendrites as described previously. Local uncaging of glutamate was performed on 5-10μm-long distal dendritic segments by three sequential trains (spaced by 2 minutes) of low frequency pulses of diffraction limited focal 405nm laser (at 0.5 Hz) at 6 different positions, which were 0.5-1 micrometers away from dendritic shafts. This protocol was shown to elicit NMDAR-mediated Ca^2+^ influx and establish prolonged sLTP at dendritic spines (*2*). All hippocampal neurons were imaged in Hibernate A Low Fluorescence Mg^2+^ free medium (BrainBits), supplemented with MNI-caged-L-glutamate at 2mM and Tetrodotoxin (TTX, Tocris) at 1.5-2μM with the temperature of a stage top incubator (Tokai Hit) set up at 35°C. Illumination takes approximately 30 seconds, and after each illumination, 11 Z-sections at a thickness of 400nm were acquired at every 10 seconds for a total duration of 600 seconds. The resulting Z-sections at each time frame were projected with maximum intensity projection feature in Fiji/ImageJ (NIH) and concatenated to construct time-lapse movies.

Glycine-mediated chemical long-term plasticity (cLTP) was used for global NMDAR stimulation as previously described (*5*). A day before the experiment, hippocampal neurons were incubated in NGM, supplemented with a selective NMDAR antagonist D-APV (Tocris) at 50μM, overnight. On the day of experiment, neurons were stimulated by briefly being incubated in Hibernate A Low Fluorescence Mg^2+^ free medium, supplemented with glycine at 200mM and Picrotoxin (Tocris) at 100μM for 5 minutes. Stimulation with glycine-containing medium was quenched and washed twice with Hibernate A Low Fluorescence (BrainBits) for following imaging. In control experiments, neurons were stimulated by cLTP protocol in the presence of D-APV. To minimize drifts, a 15-30 minute-long equilibration period right after quenching and washing was interjected before acquisition of Z-sections on multiple imaging positions ensued. Either 11 Z-sections at a thickness of 400nm or 14 Z-sections at a thickness of 300nm were acquired at 1 minute interval over 60 minutes or longer.

A widefield microscope setting, equipped with an electron multiplying CCD (EMCCD) camera (Andor), 150x 1.45 NA oil immersion objective (Olympus), automated XY stage and a piezo-Z stage (Applied Scientific Instrument) and IX81 microscope stand (Olympus), was employed for live imaging as described previously (*2*).

### smFISH and smFISH-IF of stimulated hippocampal neurons

smFISH experiments were employed for fixed imaging as described (*1*). Briefly, following cLTP stimulation, quenching and washing with NGM was performed on hippocampal neurons. For 5 minute post-cLTP time point, stimulated neurons were immediately fixed in 1x PBS (Roche), supplemented with 4% PFA (Electro Microscopy Science) and 4% sucrose (Sigma-Aldrich), 1mM MgCl_2_ (Thermo Fisher) and 0.1mM CaCl_2_ (Sigma-Aldrich), referred to as 1x PBS-MC, on ice for 20 minutes. Fixation was quenched by incubating fixed cells in 50mM glycine in 1x PBS-MC for 15 minutes at room temperature. After 3 times of wash with 1x PBS-MC, neurons were permeabilized in 1x PBS-MC, supplemented with Surfact-Amps X-100 at 0.1 % (Thermo Fisher), on ice for 15 minutes. After 3 times of wash with 1x PBS-MC, fixed and permeabilized neurons were incubated in pre-hybridization solution for smFISH, containing 10% formamide (Acros Organic) and 2x SSC (Roche) for 30 minutes at room temperature. After incubation with pre-hybridization solution, neurons were incubated in hybridization solution, consisting 10% formamide (Acros Organic), 2x SSC (Roche), 10% Dextran sulfate (Sigma-Aldrich), 2mM Ribonucleoside vanadyl complex (New England Biolabs), 1mg/ml Transfer ribonucleic acid (Sigma-Aldrich), 10U/ml SUPERaseIn RNase inhibitor (Thermo Fisher), FISH probes (Biosearch), for 4 hours at 37°C. After incubation in hybridization solution, neurons were washed in prewarmed pre-hybridization solution for 20 minutes at 37°C, and this washing step was performed twice. Lastly, neurons were washed twice with 2x SSC and twice with 1x PBS-MC before being mounted in Prolong Diamond mounting medium (Thermo Fisher). smFISH-IF was implemented with modifications of smFISH protocol, where IF staining with primary antibodies was performed simultaneously with smFISH hybridization in the same reaction buffer (i.e. hybridization solution). Fixed and permeabilized neurons were incubated in pre-hybridization solution, containing 10% formamide (Acros Organic), 2x SSC (Roche) and 0.02% BSA Fraction V (Sigma-Aldrich). And following, hybridization of neurons was performed in hybridization solution, containing FISH probes and antibodies. Following antibodies were used for smFISH-IF experiment: anti-PSD95 antibody (Antibodies Inc), anti-MAP2 antibody (PhosphoSolutions), and anti-GCN4 (SunTag) antibody (Absolute Antibody). After hybridization, neurons were briefly washed twice in BSA-containing pre-hybridization solution. Subsequently, in following 20 min-long washing steps, Alexa fluor-conjugated secondary antibodies (Invitrogen) were added at appropriate concentration (1:1000-1:1500).

A widefield microscope setting (BX63, Olympus), equipped with an CCD camera (Hamamatsu), 100x 1.40 NA oil immersion objective (Olympus) and SOLA FISH LED light engine (Lumencor), was employed for imaging smFISH and smFISH-IF samples. Randomly selected fields of views encompassing proximal and distal dendrites were acquired by Z-sectioning at a thickness of 200nm for total axial distance of 6um or 8um. Acquired Z-series were then initially inspected in Fiji/ImageJ (NIH) and selected for the best 21 Z-sections with the highest sharpness scores for downstream analyses, using Big-FISH (FISH-quant v2) python codes (*6*).

### Design and implementation of CPE-targeting morpholino ASOs

To obstruct potential interactions between *Camk2a* mRNAs and CPE-binding CPEBs via two canonical CPEs in the 3’UTR, we designed and synthesized two 28bp-long morpholino antisense oligonucleotides and 25bp-long non-targeting control ASOs with carboxyfluorescein conjugated the 3’ ends (GeneTools, Philomath, OR): (1) CPE1 ASO: 5’-TGGTCATTCAAGTTCACAGC CACAGATT-3’; (2) CPE2 ASO: 5’-ATAAACCAGACATCTCTTCTCCCCTCCC-3’; (3) Control (non-targeting) ASO: 5’-CCTCTTACCTCAGTTACAATTTATA-3’. CPE1 and CPE2 morpholino ASOs were designed to form stable complementary base pairs with portions of CPE1 and CPE2, thereby obstructing mRNA’s interaction with cognate CPEBs. To minimize possible pleotropic or indirect effects of ASO treatment, total 1uM of ASOs (CPE1 and CPE2 at 0.5uM each) per 1ml of cell culture media was transiently transfected into hippocampal neurons just 2 hours prior to cLTP protocol. Stimulated neurons were subsequently fixed at target time points and immediately proceeded to smFISH or smFISH-IF experiments as described previously.

### Image analysis of time-lapse movies

Tracking of single mRNA granule and SunTag protein signals in 2D time-lapse movies were performed as described previously and by following the instructions in the manual of TrackMate Fiji/ImageJ plugin (*7, 8*). Briefly, mRNA and SunTag protein signals in respective channels on the image of each time frame were detected by the Laplacian of Gaussian (LoG) detector with the estimated blob diameter of 5 pixels and varying thresholds. Once ideal parameters were selected, tracking was performed by the simple Linear Assignment Problem (LAP) tracker with the linking max distance and gap-closing max distance of 15 pixels and gap-closing max frame gap of 2 frames when 1 pixel was 106.7nm and frame rate was 0.0167 frame per second (i.e. 1 frame per minute). Detected spots’ spatial information (i.e. x, y coordinates) and spot-associated intensities (i.e. mean and total/integrated intensities) of each track and time frame were then extracted. Temporal aspects (e.g. decay parameter and mean lifetime) of mRNA and SunTag signals were quantified as a function of changes in intensity of signals, and the efficiency of translation of a given *Camk2a* mRNA was estimated by the frequency of *Camk2a* mRNA-SunTag ization in either dendritic shafts or dendritic spines. Comparisons between temporal changes and translation efficiencies of different experimental groups and those of control groups were made and statistical significances were computed in Prism (GraphPad), RStudio (Posit) or Jupyter Notebooks (Python).

### Image analysis of smFISH and smFISH-IF images

To isolate dendritic segments, the best 20 z-sections of randomly selected fields of views (1344×1024pixels, 86.69×66.05um) were cropped into multiple smaller fields of views (620-775×150-170pixels, 40-50×10-11um). For identification and segmentation of dendrites, Map2 IF staining for visualization of dendritic shafts or GFP IF staining for visualization of endogenously tagged GFP-CaMKIIα in both dendrites and dendritic spines was used. For detection and extraction of spatial information of single mRNA granules in dendritic shafts and spines, we utilized both radial symmetry-based mRNA detection package RS-FISH and Gaussian fitting-based mRNA detection package Big-FISH (*6, 9*). We segmented PSD95 signals in dendritic shafts and dendritic spines and GFP-CaMKIIα-filled dendritic spine heads, using Cellpose python package, and subsequently extracted their spatial information in 2D (e.g. centroids), using Big-FISH python code and Fiji/ImageJ (*10*).

Using spatial information of detected mRNA granules, PSD95 and dendritic spines, we then computed radial cross correlation of *Camk2a* mRNA signals with respect to either PSD95 or dendritic spines as a function of 2D radial distance of the pairs. Based on cross pair correlation function, where radial correlation was most enriched within 500nm distance between the pairs, we selected 500nm as an arbitrary cutoff to define association. Subsequently, we quantified the frequencies of association between PSD95 signals or spine heads with *Camk2a* mRNAs in varying conditions.

---

**Supplementary, related to Figure 1.**
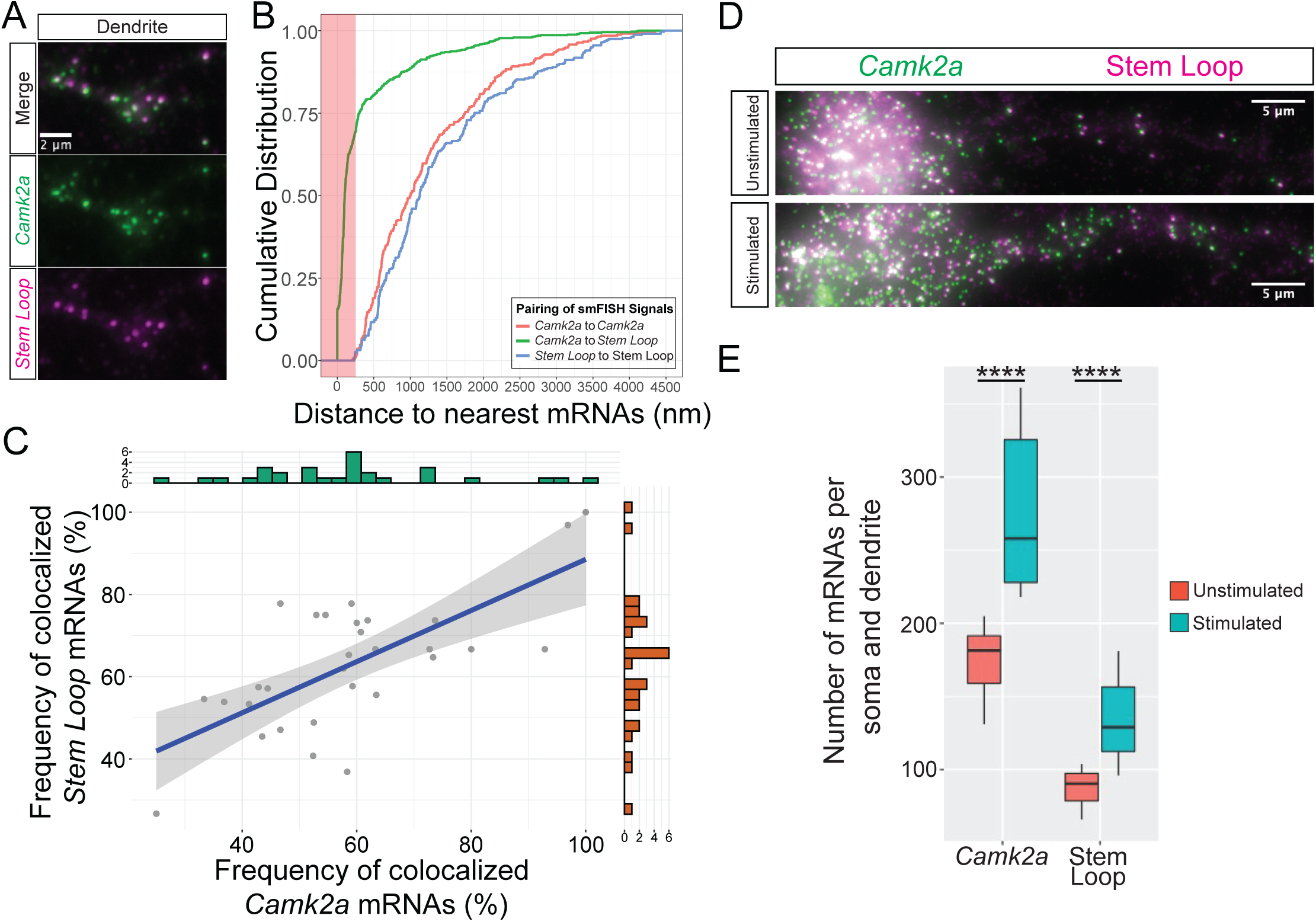
Characterization of the *Camk2a* mRNAs in the MS2/PP7 stem-loop knock-in mouse model. **(A)** Representative images of smFISH, probing for the *Camk2a* portion (CDS+UTRs, shown in green) and transgenic portion (stem loops, shown in magenta) of the endogenous mRNAs, show colocalization of the two signals in hippocampal neurons homozygous for stem-loop transgenes **(B)** Quantification shows the major fraction of *Camk2a* and Stem Loop signals are found in the proximity of one another (estimated nearest neighbor distance is ≤ 250 nm). **(C)** Correlation between co-localization frequencies of the *Camk2a* and Stem Loop smFISH signals in each analyzed neurons (n=30) is evaluated by Pearson correlation (r = 0.69; *p*-value = 2.13E-05). **(D)** Representative images smFISH, probing for both stem-loop-tagged (shown in magenta) and untagged *Camk2a* mRNAs (shown in green), show increases in abundance of the mRNAs in stimulated hippocampal neurons heterozygous for stem-loop transgenes. **(E)** Activity-induced increases in abundance of the untagged and tagged mRNAs are assessed by quantitation of mRNA molecules in proximal dendrites by one-way ANOVA, and statistical significances are indicated by horizontal bars. *p*-value annotation: * ≤0.05, **≤0.01, ***≤0.001, ****≤0.0001

**Supplementary, related to Figure 2.**
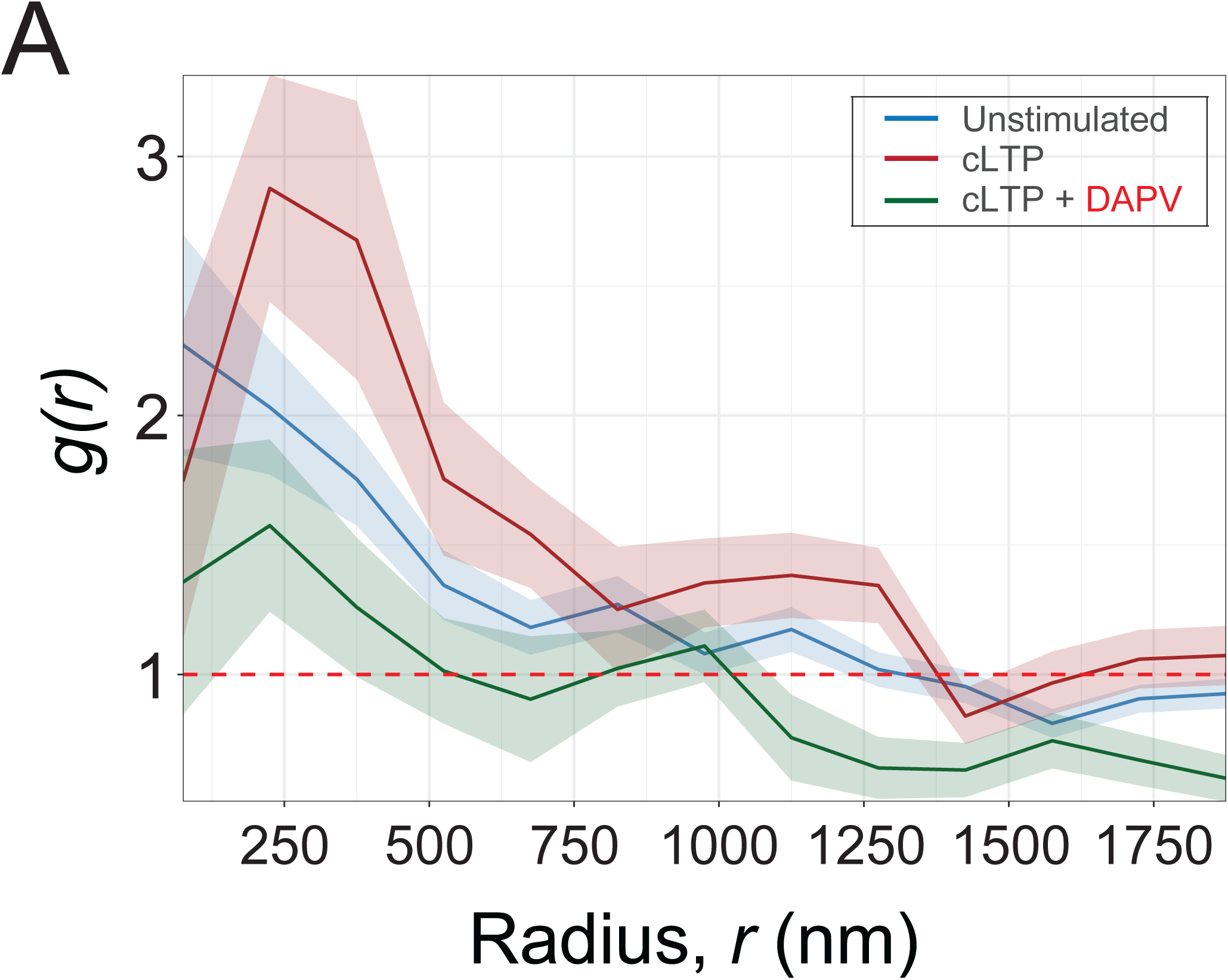
Probability of the *Camk2a* mRNAs being localized to the PSDs. **(A)** Quantification shows cross pair correlation function (*g(r)*) to find *Camk2a* mRNA with respect to centroids of PSD95 signals as a function of inter-signal distances between the pairs (radius, *r*) in unstimulated, stimulated neurons at all time points (5 min post-cLTP and 45 min post-cLTP) and stimulated by cLTP in the presence of D-APV at all time points (5 min post-cLTP + D-APV and 45 min post-cLTP + D-APV). *g(r)* >1 indicates the significant probability to find two signals at a given radius.

**Supplementary, related to Figure 3.**
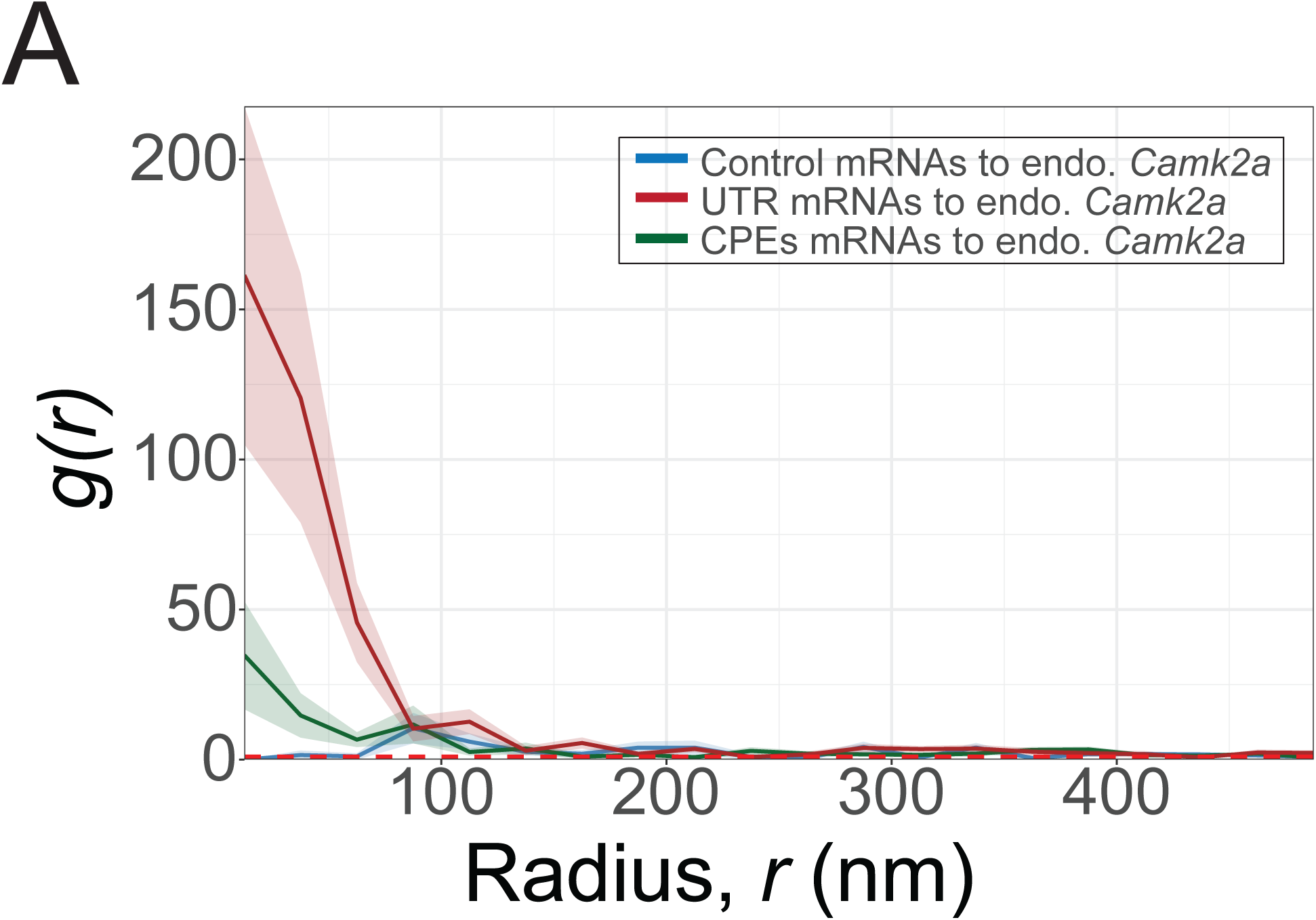
Probability of the reporter mRNAs being localized to neighboring endogenous *Camk2a* mRNAs. **(A)** Quantification shows cross correlation function (*g(r)*) to find the reporter mRNA with respect to endogenous *Camk2a* mRNA signals as a function of inter-signal distances between the pairs (radius, *r*) in stimulated neurons at 45 min post-cLTP. *g(r)* >1 indicates a significant probability to find two signals at a given radius.

**Supplementary, related to Figure 4.**
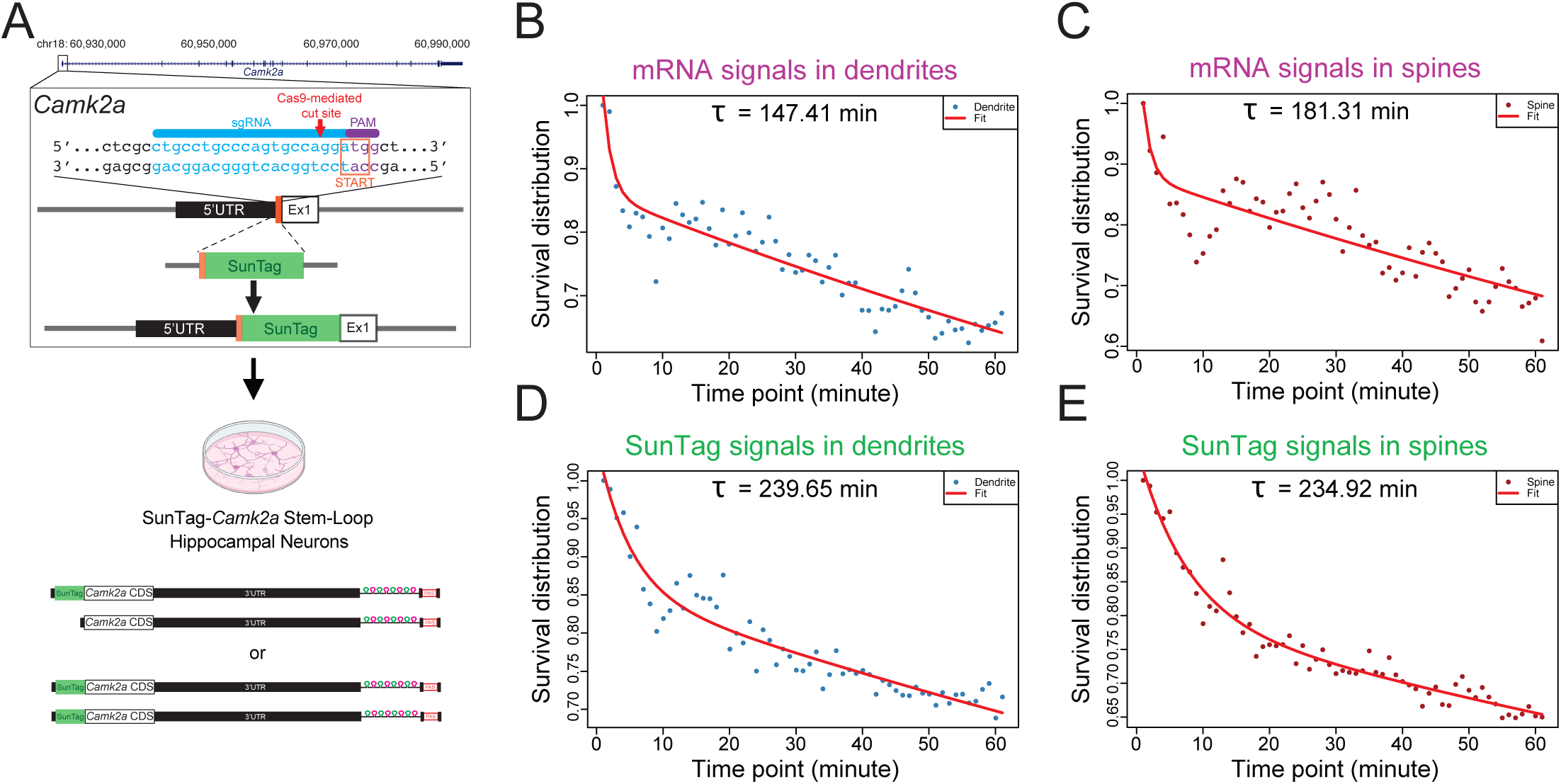
Generation of SunTag-*Camk2a* KI mouse hippocampal culture model and the lifetime estimation of the *Camk2a* mRNA and SunTag at dendritic spines in cultured neurons. (**A**) CRISPR/Cas9-mediated knock-in strategy for generating the SunTag-*Camk2a* stem-loop transgenic alleles either as homozygous or heterozygous in hippocampal neurons. (**B**) Two-component exponential decay of survival distribution of the ΔF/F of *Camk2a* mRNA signals in dendrites. (**C**) Two-component exponential decay of survival distribution of the ΔF/F of *Camk2a* mRNA signals in dendritic spines. (**D**) Two-component exponential decay of survival distribution of the ΔF/F of SunTag signals in dendrites. (**E**) Two-component exponential decay of survival distribution of the ΔF/F of SunTag signals in dendritic spines. Each data point displays the fraction of the average fluorescence intensity changes, compared to the initial intensity, in each time frame. Data points are fitted into a two-component exponential decay, indicated by red line. τ− indicates the effective lifetime, where it estimates the time that takes for a survival distribution to reach 1/e (∼0.37) of the initial fluorescence intensity. (the indicated numbers of datapoints refer to the numbers of analyzed mRNA and SunTag signals per frame from 8 time-lapse movies, n=8).

